# Polyphosphate Functions *In Vivo* as Iron Chelator and Fenton Inhibitor

**DOI:** 10.1101/2020.04.14.040345

**Authors:** Francois Beaufay, Ellen Quarles, Allison Franz, Olivia Katamanin, Wei-Yun Wholey, Ursula Jakob

## Abstract

Maintaining cellular iron homeostasis is critical for organismal survival. Whereas iron depletion negatively affects the many metabolic pathways that depend on the activity of iron-containing enzymes, any excess of iron can cause the rapid formation of highly toxic reactive oxygen species (ROS) through Fenton chemistry. Although several cellular iron chelators have been identified, little is known about if and how organisms can prevent the Fenton reaction. By studying the effects of cisplatin, a commonly used anticancer drug and effective antimicrobial, we discovered that cisplatin elicits severe iron stress and oxidative DNA damage in bacteria. We found that both of these effects are successfully prevented by polyphosphate (polyP), an abundant polymer consisting solely of covalently linked inorganic phosphates. Subsequent *in vitro* and *in vivo* studies revealed that polyP provides a crucial iron reservoir under non-stress conditions, and effectively complexes free iron and blocks ROS formation during iron stress. These results demonstrate that polyP, a universally conserved biomolecule, plays a hitherto unrecognized role as an iron chelator and an inhibitor of the Fenton reaction.

## Introduction

Cisplatin (cis-diaminedichloroplatinum (II)), originally identified as an inducer of bacterial filamentation (Rosenberg et al., 1965), is one of the most widely used drugs in cancer treatment (Galanski et al., 2005, Hannon, 2007). Early mechanistic studies suggested that cisplatin elicits cytotoxicity by acting as a DNA-damaging agent, preferentially crosslinking neighboring purines (Bancroft et al., 1990, Eastman, 1987, Sherman & Lippard, 1987). More recent studies, however, revealed that cisplatin also induces cell death in denucleated cells by causing mitochondrial and endoplasmic reticulum stress (Mandic et al., 2003, Yang et al., 2006). Indeed, only a small fraction of the intracellular cisplatin pool appears to reach the nucleus, whereas the vast majority binds to the sulfur-containing side chains in proteins as well as to thiol-containing compounds (Ishikawa & Ali-Osman, 1993, Karasawa et al., 2013, Lin et al., 2002, Peleg-Shulman et al., 2002). Studies in mouse models and ovarian cancer cell lines revealed that tumorous cells gain resistance against cisplatin by increasing their levels of cysteine-enriched peptides (i.e., glutathione) and proteins (i.e., metallothioneins), which capture cisplatin before it reaches the DNA (Holford et al., 2000, Kelland, 2007, Kelley et al., 1988). This cellular response also seems to aid in mitigating oxidative stress, a frequently observed side effect of cisplatin treatment (Santos et al., 2007).

Recent studies from our lab revealed that upon cisplatin treatment, cancer cells drastically upregulate and redistribute their levels of inorganic polyphosphate (polyP) (Xie et al., 2019). PolyP is a polymer of up to 1000 inorganic phosphate (Pi) molecules, linked by high-energy phospho-anhydride bonds (reviewed in (Xie & Jakob, 2019)). In bacteria, polyP protects against a variety of different stress conditions (i.e., oxidative stress, heat stress), stimulates biofilm formation, and regulates virulence (Rao et al., 2009) (Gray et al., 2014). Some of these functions can be explained by the ability of polyP to work as a protein-stabilizing scaffold (Gray et al., 2014). As such, polyP protects soluble proteins against stress-induced aggregation while promoting the formation of functional amyloids, including those involved in biofilms (Cremers et al., 2016). Other potential functions that have been associated with polyP are based on its chemical features as a buffer or high-energy storage molecule (reviewed in (Gray & Jakob, 2015)).

To gain more insights into the working mechanism of cisplatin and the role that polyP might play in the cellular response to this drug, we compared the effects of cisplatin treatment on wild-type and polyP-deficient *E. coli*. Our studies demonstrated that cisplatin triggers a gene expression pattern in wild-type bacteria that is consistent with the inactivation of the repressor Fur, the master regulator of iron homeostasis (Seo et al., 2014). The resulting gene expression changes lead to an apparent increase in iron uptake, whose deleterious effect is effectively mitigated by endogenous polyP. Deletion of the polyP synthesizing machinery causes a dramatic increase in cisplatin-induced mutagenesis rate and cell death. Both phenotypes are fully prevented in polyP-depleted bacteria by overexpressing iron storage proteins or by globally reducing the number of iron-containing proteins. Subsequent *in vivo* and *in vitro* studies revealed that polyP acts as a hitherto unknown iron-storage molecule under both stress and non-stress conditions, and, by chelating labile iron, acts as a physiologically relevant Fenton inhibitor.

## Result

### PolyP protects *E. coli* against cisplatin toxicity

Despite its prevalent use in anti-tumor treatment and its known broad antibacterial activity, the exact mechanism by which cisplatin kills cells is still not fully understood. To obtain more detailed insight into the cellular effects of cisplatin, we investigated the responses to and defenses against cisplatin toxicity in bacteria. Based on our recent discovery that polyP serves as an active defense mechanism against oxidative protein damage in bacteria (Gray et al., 2014), we compared the cisplatin-sensitivity of wild-type *E. coli* with mutant strains that lack either the polyP-synthesizing polyP kinase PPK (polyP-deplete) or the polyP-hydrolyzing enzyme exopolyphosphatase PPX (polyP-replete). We grew all three strains to mid-log phase in minimal MOPS glucose (MOPS-G) medium and exposed them to increasing amounts of cisplatin either on plates (Figure 1A) or in liquid culture (Figure S1A). Whereas *E. coli* wild-type or the *ppx* deletion mutant strain did not show any growth defects when incubated on plates supplemented with 4 μg/ml of cisplatin, the *ppk* deletion mutant, which lacks detectable polyP levels (Gray et al., 2014), showed a reproducible three to four log reduction in cell survival (Figure 1A). We obtained very similar results when we conducted the experiments in liquid culture. After a 20h treatment with 10 μg/ml cisplatin in liquid media, wild-type *E. coli* showed an about 3-log decrease in survival while the *ppk* deletion strain showed a greater than 6-log decrease (Figure S1A).

**Figure 1.**
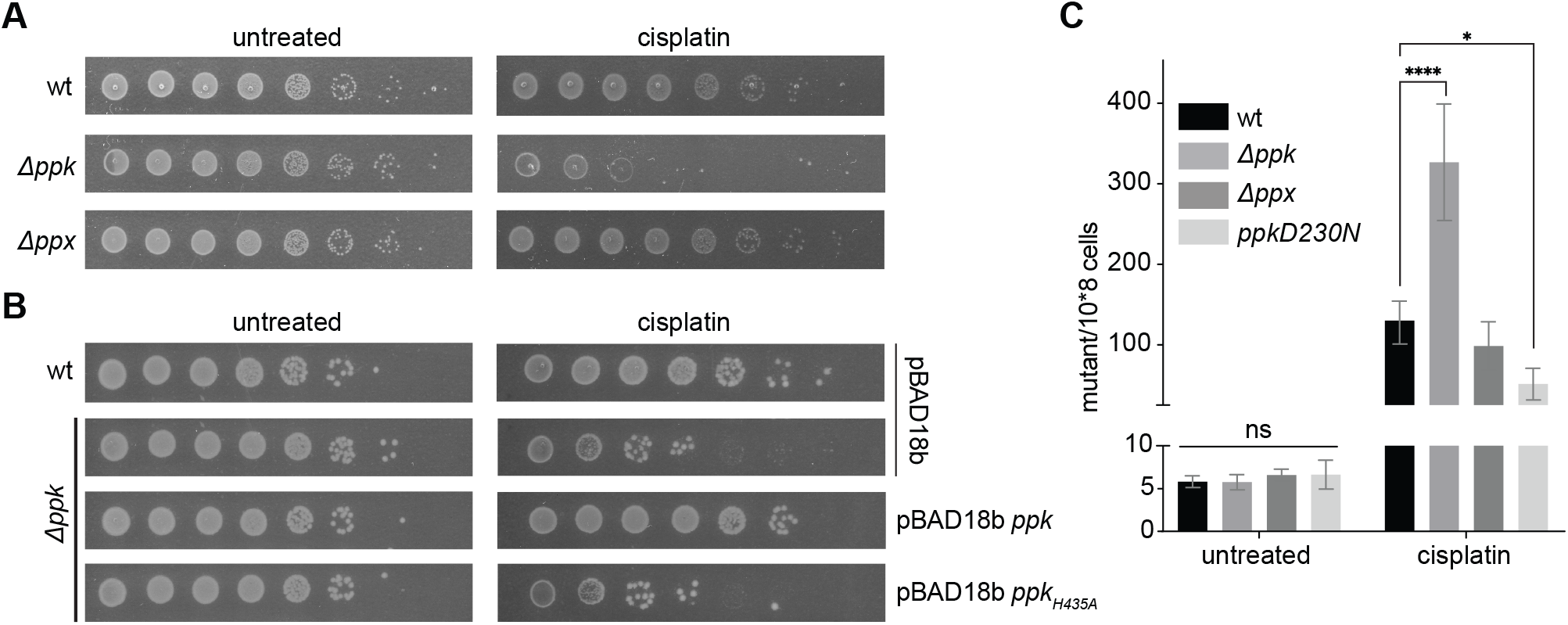
PolyP confers cisplatin resistance in *E. coli*. **A.** Logarithmically growing *E. coli* MG1655 wild-type (wt), *Δppx* or *Δppx* were 10-fold serially diluted, spotted onto M9-G plates containing 4 μg/ml cisplatin and incubated at 37°C overnight. **B.** Logarithmically growing *E. coli* wild-type containing the empty vector pBAD18b, or the *ppk* mutant strain containing either the empty vector or expressing wild-type PPK or the enzymatically inactive PPK_H435A_ mutant were 10-fold serially diluted, spotted onto plates containing 0.02% w/v arabinose and 4 μg/ml cisplatin, and incubated at 37°C overnight. Experiments shown in A, B were conducted at least 4 times, and a representative result is shown. **C.** The mutagenesis rates of logarithmically growing wild-type (wt), *Δppx*, *ΔppK* or *ΔppK* expressing the highly active PPK-D230N mutant protein before and 1h after treatment with 10 μg/ml cisplatin in liquid culture was determined by counting the number of colonies able to grow on rifampicin plates (n=3; *, P<0.05; ****, P<0.0001; ns, non-significant, one-way ANOVA).

Analysis of the endogenous polyP levels in wild-type *E. coli* did not reveal any significant upregulation of polyP in response to cisplatin treatment (Figure S1B), suggesting that the steady-state levels of polyP that are present in stationary phase *E. coli* (Rao & Kornberg, 1996) are sufficient to confer the observed protection. To ascertain, however, that polyP synthesis is indeed required for the observed cisplatin resistance in wild-type *E. coli*, we transformed the *ppk* deletion strain with plasmids encoding for either the native PPK protein or the previously characterized, catalytically inactive variant PPK-H435A (Kumble et al., 1996). As shown in Figure 1B, the expression of wild-type PPK fully rescued the cisplatin-sensitivity of the *ppk* deletion strain. The expression of the catalytically inactive PPK variant, on the other hand, failed to rescue the growth defect. These results strongly suggested that endogenous levels of polyP are necessary and sufficient to protect *E. coli* against chronic cisplatin stress.

### PolyP protects *E. coli* against cisplatin-induced DNA damage

To begin to understand how polyP protects bacteria against cisplatin toxicity, we analyzed the expression of select heat shock and SOS-response genes in cisplatin-treated wild-type, *ppk* and *ppx* deletion mutants. This line of experiments was instigated by our previous study, which showed that during severe oxidative stress, polyP-deficient bacteria upregulate their heat shock gene expression levels in an apparent attempt to compensate for the lack of polyP’s chaperone function (Gray et al., 2014). Analysis of the mRNA levels of IbpA and DnaK, two genes whose expression levels are highly responsive towards protein unfolding stress in *E. coli* (Guisbert et al., 2008), did not reveal any significant change upon cisplatin treatment in any of the three tested strains (Figure S1C). In contrast, however, we found that the mRNA levels of the gene encoding for the cell division inhibitor SulA, a major component of the SOS response and an inducer of bacterial filamentation, significantly increased upon cisplatin treatment in the wild-type *E. coli* and *ppx* deletion strain, and went up even more in bacteria lacking polyP. These results agreed with the original observation that cisplatin treatment triggers bacterial filamentation (Rosenberg et al., 1965), and indicated that at the concentrations used, cisplatin works as a DNA rather than a protein-damaging reagent in bacteria.

To assess the levels of DNA damage that cisplatin elicits in the *ppk* deletion strain *versus* in bacteria that contain measurable levels of polyP, we determined the mutagenesis rates before and after cisplatin-treatment by counting the number of bacteria that gain the ability to grow on rifampicin plates (Touati et al., 1995). Rifampicin resistant mutations in the RNA polymerase gene *rpoB* arise from single base substitutions, which prevent rifampicin from binding to and hence inhibiting RNA polymerase (Campbell et al., 2001). Under non-stress condition, the spontaneous rate of mutagenesis in all three tested strains was similarly low (< 10 mutants per 10^8^ cells) (Fig. 1C). Not unexpectedly given cisplatin’s mode of action, this number drastically increased upon cisplatin treatment. However, the mutagenesis rate of the *ppk* deletion was almost 3-fold higher compared to the mutagenesis rates of cisplatin-treated wild-type *E. coli* or the *ppx* deletion strain. Expression of a highly active PPK variant PPK-D230N, which substantially increases the steady state levels of polyP *in vivo* (Rudat et al., 2018), reduced the mutagenesis rate of the *ppk* deletion strain to levels that were even lower than the ones observed in cisplatin-treated wild-type *E. coli* (Fig. 1C). These results suggested a previously unrecognized function of polyP in bacterial DNA damage control.

### Cisplatin triggers extensive transcriptional changes in wild-type and polyP-depleted *E. coli*

We conducted RNAseq analysis to compare the transcriptional profile of wild-type and *Δppk* cells in response to a sub-lethal dose of cisplatin (i.e., 20 μg/ml cisplatin for 15 minutes). Compared to the respective untreated controls, we identified more than 1,100 differentially expressed genes (DEGs) in each strain (see supplemental Tables S1-S4). We categorized the DEGs into clusters of ontology according to their functional annotation (GOterm) (Ashburner et al., 2000), and focused our primary analysis on the two following groups: 1) DEGs in wild-type *E. coli* upon cisplatin treatment (Figure 2A; Tables S1, S4) since we reasoned that those genes will likely reveal the cellular effects that cisplatin exerts in bacteria; and 2) DEGs in cisplatin-treated wild-type *versus Δppk* strain (Figure 2B; Tables S3, S4) since we deduced that those genes will likely provide mechanistic insights into how polyP protects bacteria against cisplatin stress. The most upregulated genes in cisplatin-treated wild-type *E. coli* compared to the untreated control belonged to members of the SOS response (Figure 2A, Table S1). This was not an unexpected result since the SOS response is the canonical response to DNA damage. It controls more than 40 genes involved in transcriptional regulation, DNA repair mechanism, cell cycle arrest, and error-prone DNA synthesis (Courcelle et al., 2001, Friedberg, 1985, Friedberg et al., 1995) (Figure 2A). Many of the next most significantly altered cluster of DEGs in cisplatin-treated wild-type *E. coli* included genes involved in iron-sulfur cluster assembly, iron uptake, enterobactin synthesis and iron regulation (Fig. 2A, red bars). These results suggested that cisplatin treatment causes a disturbance in the cellular iron homeostasis. Indeed, when we compared the DEGs in cisplatin-treated wild-type with the DEGs identified under iron starvation or repletion conditions (Seo et al., 2014), we observed a statistically highly significant overlap (Fig. 2C). Particularly striking was the overlap between genes differentially regulated by cisplatin and genes previously reported to be directly controlled by the master transcriptional repressor *F*erric *U*ptake *R*egulator FUR (Seo et al., 2014) (Figure 2C). In fact, the expression pattern in cisplatin-treated wild-type bacteria strongly resembled the expression pattern in *E. coli* mutants lacking functional FUR, i.e., the up-regulation of genes involved in iron uptake and the selective down-regulation of genes involved in iron utilization (Figure 2D). These results suggested that cisplatin treatment affects iron homeostasis in wild-type bacteria, either by triggering or signaling an iron starvation response. Other significantly enriched gene clusters in cisplatin-treated wild type bacteria included genes involved in fermentation and anaerobic ETC, as well as nitrate assimilation, sulfur and phosphate metabolism (Figure 2A).

**Figure 2.**
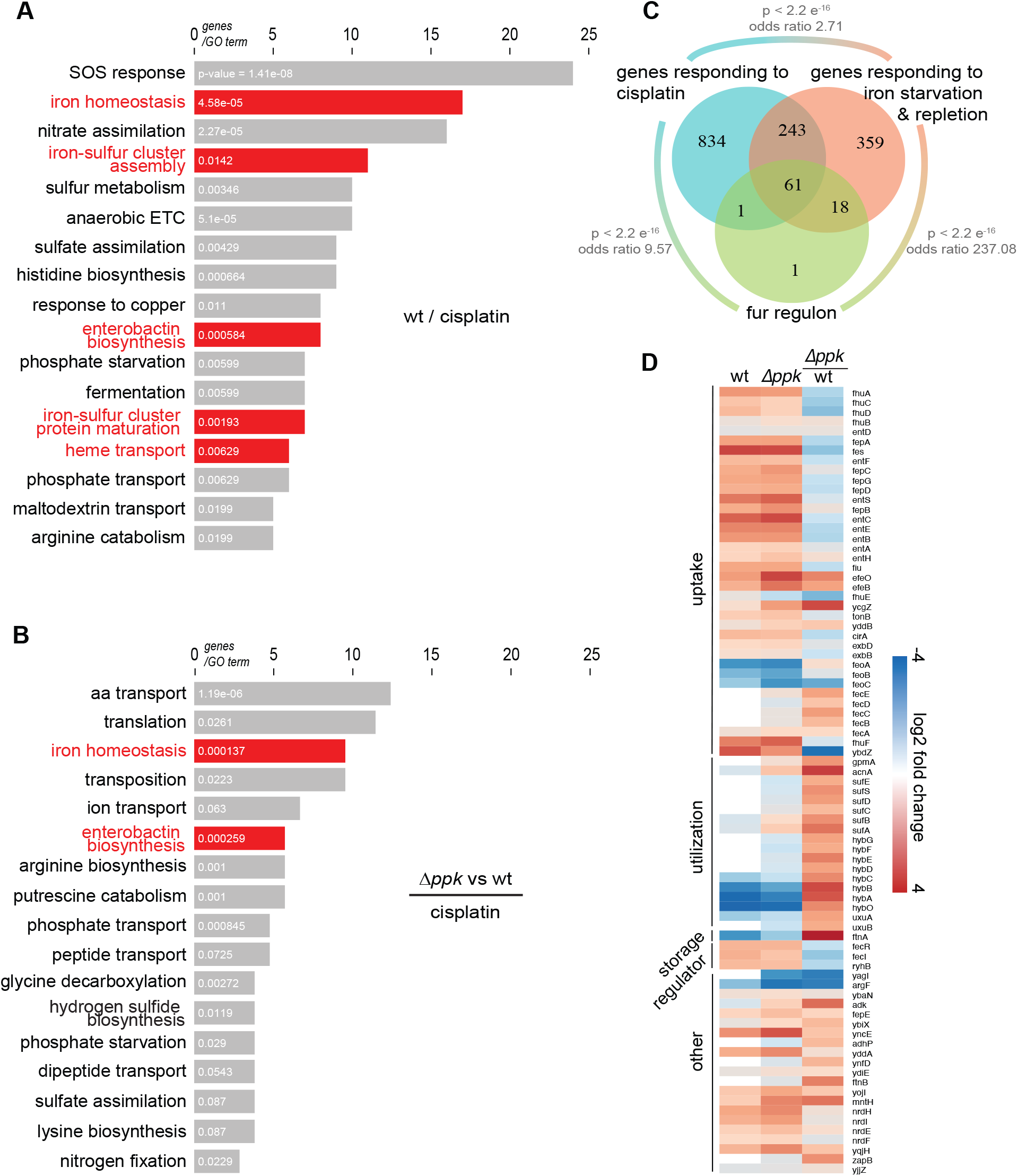
Global gene expression changes in response to cisplatin treatment. **A.** Functional classification (GOterm) of genes differentially expressed in wild-type *E. coli* upon 15 min treatment with a non-lethal cisplatin concentration (20 μg/ml) (Table S1, S4). Categories related to iron metabolism are highlighted in red. **B.** Functional classification (GOterm) of genes differentially expressed in *Δppk* and wild-type *E. coli* upon 15 min treatment with a non-lethal cisplatin concentration (20 μg/ml) (Table S3, S4). *P*-values are shown in white over each bar, from a modified Fisher’s Exact test. **C.** Venn diagram of genes belonging to the *fur* regulon (green circle), genes differentially expressed in wild-type *E. coli* upon cisplatin treatment (cyan circle), and genes differentially expressed in a *fur* deletion strain under iron repletion and starvation conditions (Seo et al., 2014) (red circle). *P*-values and odds ratios from Fisher Exact Tests. **D.** Heatmap of Fur-regulated genes, which are differentially expressed in cisplatin treated wild-type (wt) and *Δppk* cells. Data are log2 fold change, range: −4 (blue) to 4 (red). Ratio of *ppk* to wild-type is shown in the right column. Genes are organized according to their functional annotations.

### Cisplatin elicits iron stress in polyP-deficient *E. coli*

Direct comparison of the DEGs in cisplatin-treated wild-type *E. coli* and polyP-deficient mutant cells revealed numerous gene clusters that responded similarly to cisplatin treatment in both strains (compare Fig. 2A with Fig. S2A; Table S3). In stark contrast, however, we found dramatic differences between the two strains in the expression of genes involved in amino acid transport, translation, transposition, and, particularly, iron homeostasis (Fig. 2B). Given the close connection between iron, oxidative stress, and DNA damage, we subsequently focused on the DEGs associated with iron homeostasis. We observed that polyP-depleted cells respond to cisplatin treatment with a much less pronounced expression of iron uptake genes and an even more pronounced downregulation of iron utilization genes (Figure 2D, Table S3). This expression pattern suggested that polyP-depleted bacteria experience a relative increase in the intracellular labile iron pool upon cisplatin treatment, which would explain their relative decrease in survival and increase in mutagenesis rate compared to wild-type bacteria (Fig. 1A, C). Given the polyanionic structure of polyP, and its previously shown ability to interact with divalent metals, such as Ca^2+^, Mg^2+^ and certain heavy metals (Keasling, 1997, Ruiz et al., 2011), we therefore considered the possibility that polyP serves as a hitherto unknown iron chelator. PolyP might complex excess iron as it is being taken up from the extracellular space and/or released from iron-containing proteins during cisplatin treatment. We reasoned that if this model were to be correct, we should be able to specifically rescue the cisplatin-sensitivity of the *ppk* deletion strain by reducing the intracellular iron load during cisplatin treatment. To test this idea, we devised three different strategies; i) decreasing the extracellular iron concentration, which should reduce the amount of Fe-uptake during cisplatin stress (Braun, 2001); ii) overexpressing the iron storage protein FtnA, which should compensate for the lack of polyP; or iii) overexpressing the small RNA *ryhB*, which promotes the degradation of numerous mRNAs coding for iron-containing proteins, and hence reduces the number of iron-containing proteins in the cell (Massé & Gottesman, 2002). For our first strategy, we grew wild-type and *ppk* deletion strains in liquid medium supplemented with either normal (+ Fe) or low (-Fe) concentrations, exposed them to our previously established cisplatin treatment, and determined cell survival and mutagenesis rates. The result that we obtained was fully consistent with our hypothesis; lowering the iron concentration in the media dramatically increased the survival and reduced the mutagenesis rates of the *ppk* deletion strain to levels directly comparable to cisplatin-treated *E. coli* wild-type (Figure 3A, B). In contrast, cisplatin treatment under low *versus* normal iron conditions did not yield any noticeable difference in the cisplatin resistance of wild-type *E. coli.* These results suggested that iron taken up from the media in response to cisplatin treatment was successfully complexed or otherwise neutralized by polyP. We obtained a very similar result upon overexpression of the plasmid-encoded iron storage protein FtnA, which significantly improved survival and reduced the mutagenesis rates in the cisplatin-treated *ppk* deletion strain but not in wild-type *E. coli* (Fig. 3C, D). These results demonstrated that expression of a protein-based iron chelator fully compensates for the lack of polyP. We finally tested the effect of *ryhB* overexpression, a strategy that bacteria typically use to reduce the number of Fe-containing proteins under low-iron conditions (Massé & Gottesman, 2002). Indeed, and in agreement with our previous results, we found that the overexpression of *rhyB* restored the cisplatin resistance of polyP-depleted cells to wild-type like levels while rendering the resistance of wild-type *E. coli* unaltered (Figure 3E, F). These results suggested that iron complexed in metalloproteins might contribute to the iron toxicity in the absence of polyP. Since previous reports documented that cisplatin covalently binds cysteine residues in proteins (Ishikawa & Ali-Osman, 1993, Kelland, 2007, Williams et al., 2004), including those involved in iron binding (Bischin et al., 2011), we finally tested the idea that cisplatin targets iron-containing proteins in bacterial cell lysates. We therefore prepared wild-type or *Δppk* crude extracts, and measured the activity of aconitase, an enzyme, whose iron-sulfur cluster is coordinated via three oxidation-sensitive cysteines. We found that increasing concentrations of cisplatin indeed increasingly reduced the activity of aconitase at similar level in both wild-type and *Δppk* extracts (Figure S3). Together, these results suggested that endogenous polyP provides an effective non-proteinogenic mechanism to protect bacteria against cisplatin-mediated accumulation of free iron and Fe-mediated oxidative damage.

**Figure 3.**
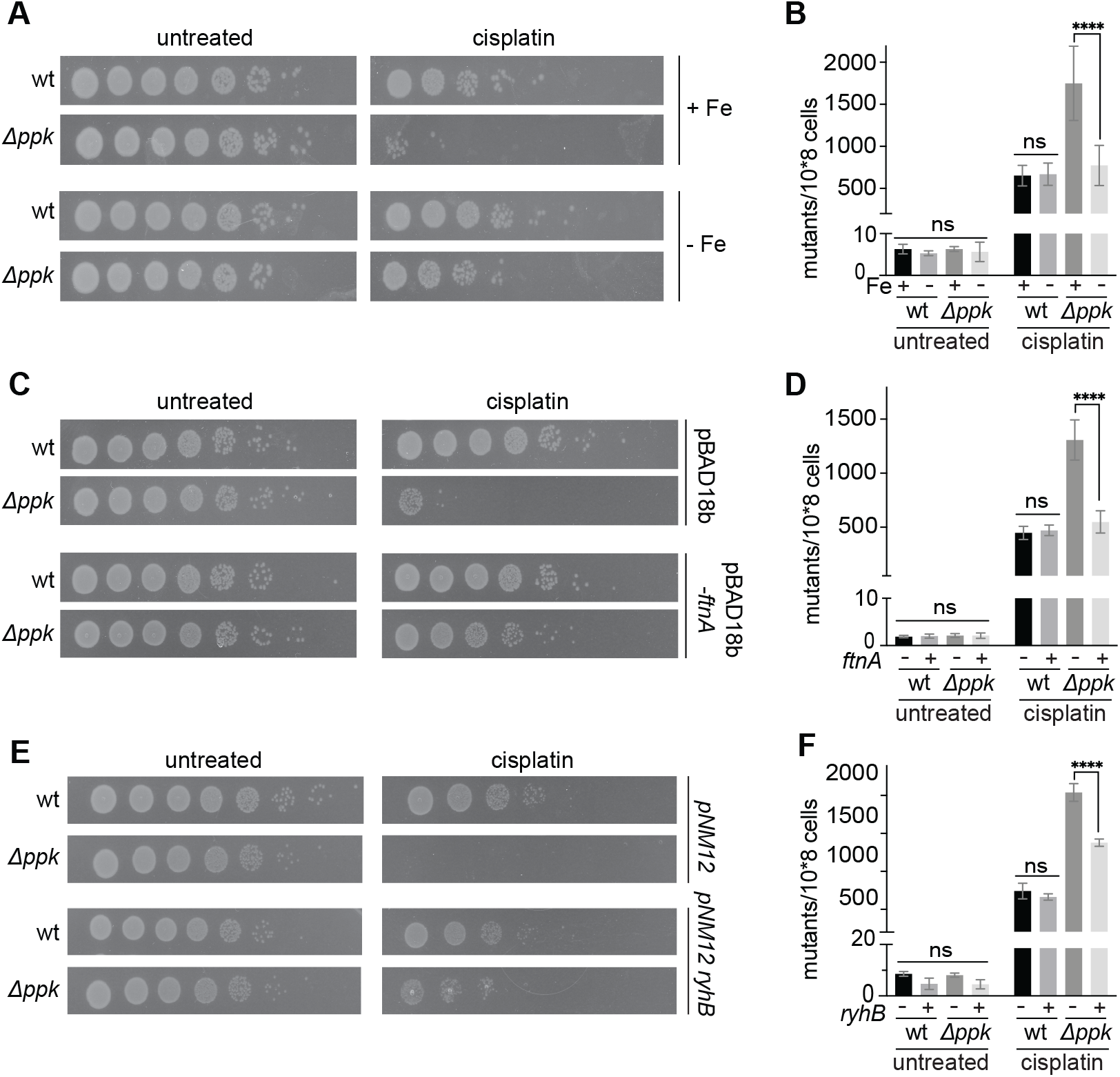
Low-iron conditions reduce cisplatin toxicity in polyP deficient *E. coli*. **A, B.** Exponentially growing *E. coli* wild-type (wt) and *ppk* deletion strains were exposed to 10 μg/ml cisplatin in untreated (+Fe) or chelexed (-Fe) M9-G media for (A) 20 h to determine survival or (B) for 1 h to determine the rate of mutagenesis. **C, D.** Exponentially growing *E. coli* wild-type and *ppk* deletion strains carrying either an empty plasmid or a plasmid overexpressing the iron storage protein *ftnA* were exposed to 10 μg/ml cisplatin in M9-G media for either 20 h to determine survival (C) or for 1 h to determine the rate of mutagenesis (D). **E, F**. Exponentially growing wild-type and *Δppk* cells carrying an empty plasmid or a plasmid overexpressing *ryhB* were serially diluted, spotted onto plates containing 0.02% w/v arabinose and 4 μg/ml cisplatin and incubated at 37°C to determine survival (E), or exposed to 10 μg/ml cisplatin in M9-G media for 1h to determine the rate of mutagenesis (F). n=3; ****, P<0.0001; ns, non-significant; one-way ANOVA.

### PolyP acts as a physiologically relevant iron-storage molecule *in vivo*

Iron homeostasis is a tightly regulated mechanism, necessitated by the fact that unbound labile iron is highly toxic under aerobic growth conditions. This toxicity appears to be primarily caused by the ability of free iron to interact with peroxide (i.e., Fenton reaction), which leads to the production of highly reactive hydroxyl radicals (Dixon & Stockwell, 2014). To further evaluate the idea that polyP serves as a general, hitherto unrecognized iron storage molecule in bacteria, we turned to a mutant strain of *E. coli*, which lacks the master iron repressor Fur. Deletion of Fur triggers an iron starvation response, which, similar to the situation we observed in cisplatin-treated wild-type *E.coli*, leads to gene expression changes that are geared towards replenishing the intracellular iron pool (Andrews et al., 2003, Massé & Gottesman, 2002). As a cellular consequence, *fur* deletion strains suffer from an intracellular iron overload, which causes severe growth defects and significantly higher mutagenesis rates under aerobic but not under anaerobic growth conditions (Touati et al., 1995). To test whether polyP functions as an iron storage molecule in a *Δfur* deletion strain, we generated *ΔfurΔppk* double deletion strains, and analyzed growth and mutagenesis rates under both aerobic and anaerobic growth conditions. As shown in Figures 4A-C, additional deletion of *ppk* in the *Δfur* deletion strain significantly aggravated the growth defect and increased the mutagenesis rate specifically under aerobic growth conditions (Fig. 4A-C). These results were fully consistent with our prior observations and supported our conclusion that polyP protects bacteria generally against conditions of Fe-overload. To finally test whether polyP also serves as an iron reservoir under non-stress conditions, we cultivated *E. coli* wild type, *Δppk* and *Δppx* strains in M9 minimal medium in the presence of increasing concentrations of the iron chelator 2,2’-dipyridyl, using gluconate as sole carbon source. By offering solely gluconate, bacterial growth depends on the activity of the iron-sulfur cluster protein gluconate dehydratase, and hence on the availability of intracellular iron (Gardner & Fridovich, 1991). Addition of 2,2’-dipyridyl to the growth media reduces the intracellular iron pools and, at sufficiently high levels, prevents bacterial growth as it depletes the Fe-S cluster in gluconate dehydratase. As shown in Fig. 4D (and Fig. S4A, B), whereas the *Δppx* strain was slightly more resistant towards the presence of the chelator compared to wild-type *E. coli*, the *Δppk* strain was significantly more sensitive. Less than 100 μM 2,2’-dipyridyl in the growth media was sufficient to decrease the relative growth rate of the *Δppk* strain by 50% while more than 150 μM of the chelator was necessary to trigger the same growth defect in wild-type *E. coli* (Fig. 4D, Figs. S4A, B). These results demonstrated that lack of endogenous polyP drastically increases the sensitivity of *E. coli* towards the presence of iron chelators in the media and supports the conclusion that polyP serves as a physiologically relevant iron storage molecule in *E. coli.*

**Figure 4.**
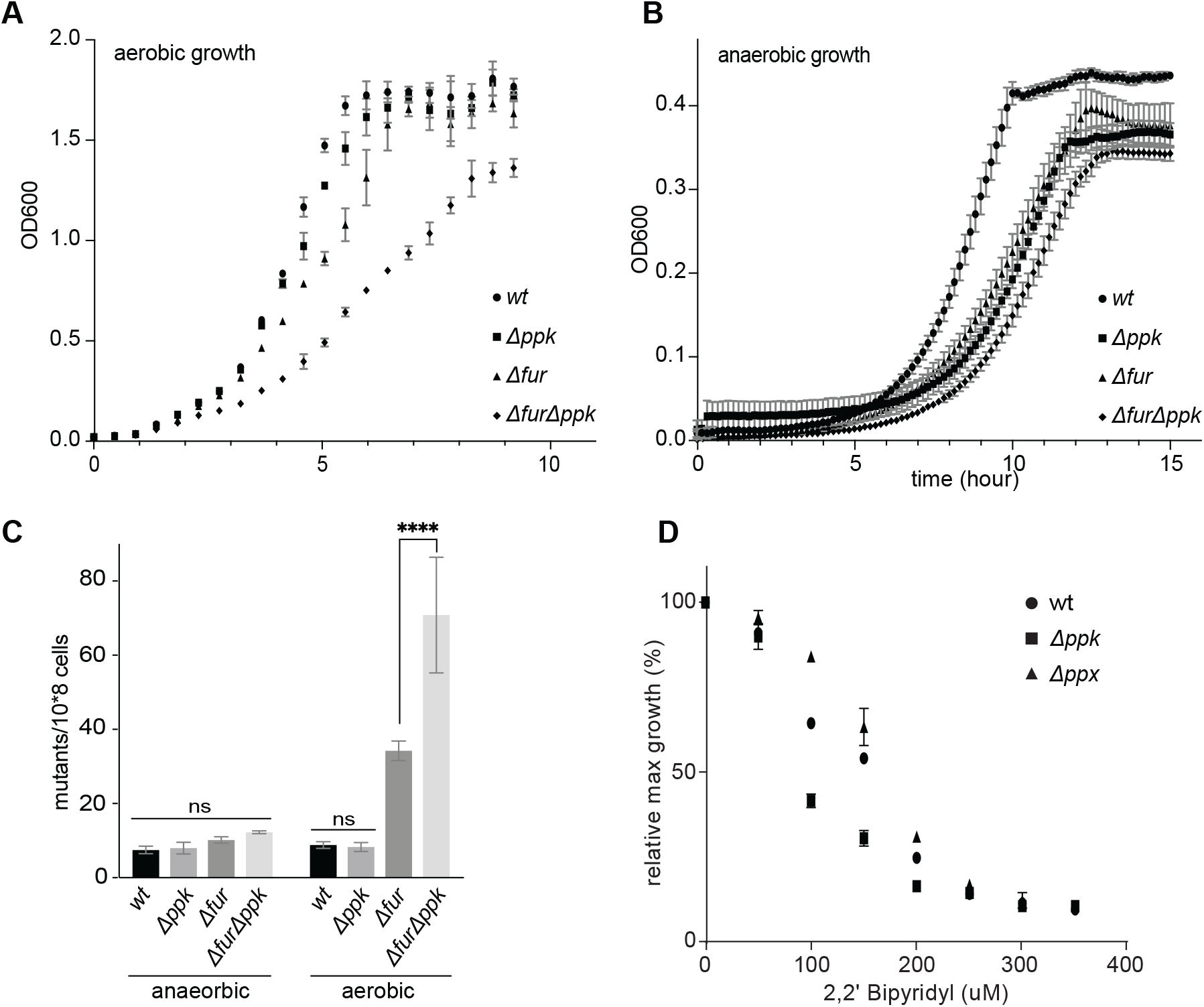
PolyP acts as an iron-storage molecule *in vivo*. **A, B.** Growth of *E. coli* wild-type, *Δppk*, *Δfur* and *ΔfurΔppk* strains in MOPS-G media under aerobic (A) or anaerobic condition (B) conditions. **C.** Mutagenesis rate of each strain under aerobic and anaerobic growth conditions. (n=3; ****, P<0.0001; ns, non-significant; one-way ANOVA). **D.** Exponentially growing *E. coli* wild-type, *Δppk* and *Δppx* deletion strains were diluted into M9-Gluconate supplemented with increasing concentrations of the iron chelator 2,2’ bipyridyl. The maximal growth achieved after 18h incubation at 37°C was normalized against the growth in M9-Gluconate without chelator. Error bars are ± one standard deviation for technical triplicates. Experiments were performed in triplicates (for biological replicates see Fig. S4A, B) and a representative result is shown.

### PolyP protects against Fe-mediated DNA damage *in vitro*

To directly test whether polyP, through complexing iron, mitigates the Fenton reaction, we monitored the H_2_O_2_ / FeSO_4_ -mediated oxidation of 2,2’-azino-bis(3-ethylbenzothiazoline-6-sulphonic acid) (ABTS) (Zheng & Huang, 2014) in the presence of increasing amounts of polyP (Fig. 5A). ABTS, once oxidized by H_2_O_2_ / FeSO_4_ -produced hydroxyl radicals, shows a strong absorbance signal at 414 nm. As shown in Fig. 5A, the presence of polyP prevented ABTS oxidation in a concentration-dependent manner, indicating that chelation of Fe^2+^ by polyP inhibits the Fenton reaction. In fact, 20 μM polyP_300_ (in P_i_ units) was sufficient to prevent oxidation of ABTS by a mixture of 5 μM FeSO_4_ and 20 μM H_2_O_2_ (Fig. 5A). To test whether the association of Fe^2+^ with polyP also protects DNA against Fe-mediated oxidative damage, we incubated 10 μM of linearized DNA with a mixture of 50 μM Fe^2+^ / 5 mM H_2_O_2_ in the absence and presence of polyP_300_ (Fig. 5B). Whereas in the absence of polyP, all of the DNA was oxidatively degraded within a 30 min incubation period, the presence of 5 mM polyP_300_ (in Pi-units), almost completely prevented the degradation of DNA. Presence of inorganic phosphate (Pi) in the form of either sodium- or potassium phosphate did not have any protective effect even when used at concentrations as high as 75 mM (Fig. 5B), confirming that the polyanionic nature of the polyP chain is necessary to sequester iron in a non-reactive state. In summary, these results demonstrate that polyP acts as a physiologically relevant iron storage molecule, capable of preventing the production of hydroxyl radical by the Fenton reaction.

**Figure 5.**
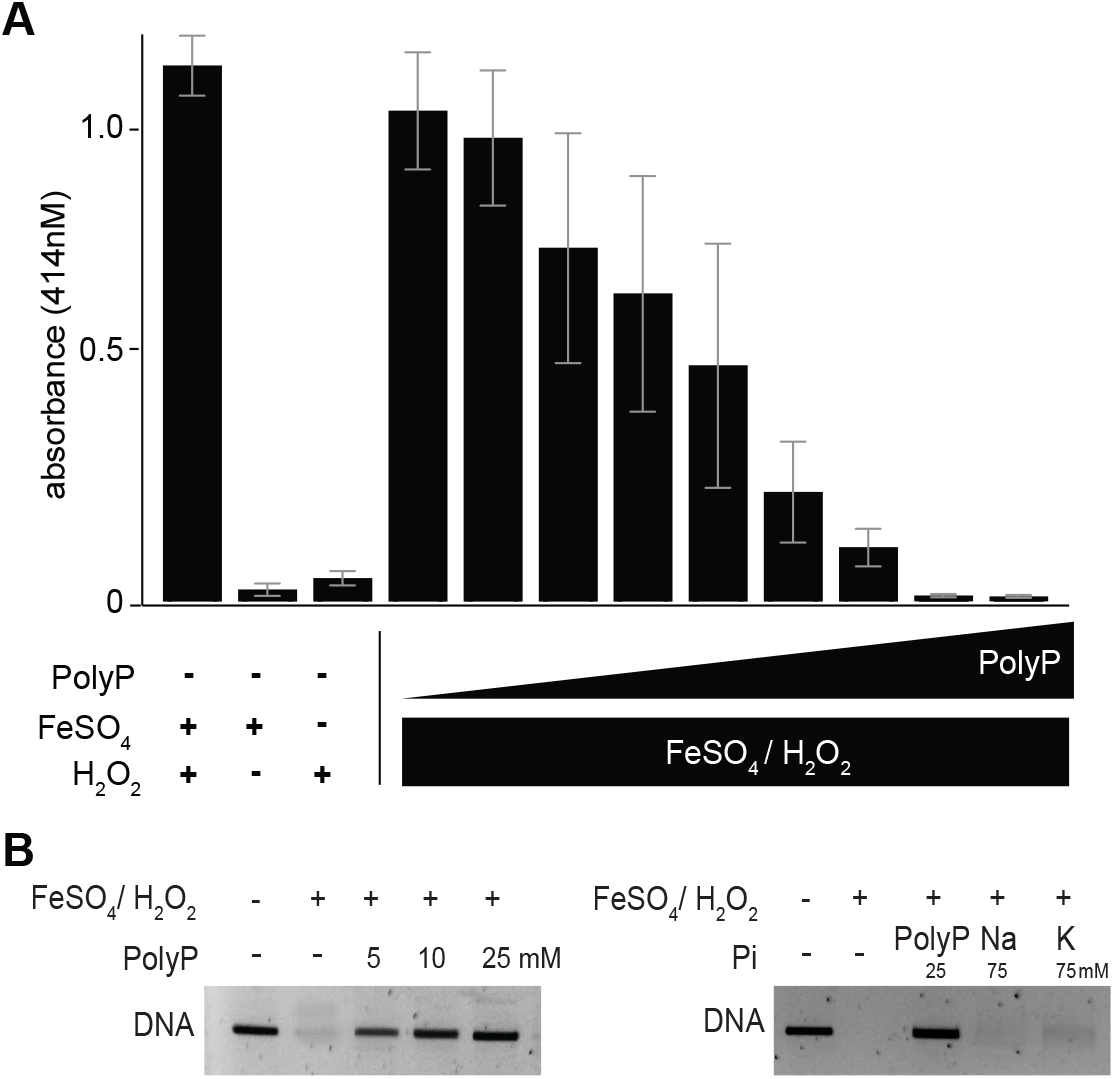
PolyP inhibits Fenton reaction *in vitro*. **A**. ABTS oxidation by 20 μM H_2_O_2_ and 5 μM FeSO_4_ was monitored at 414 nm in the absence or presence of 1, 2, 3, 4, 5, 7.5, 10, 20 or 30 μM of polyP_300_ (in Pi units). No significant oxidation was observed in the presence of H_2_O_2_ or FeSO_4_ alone (n= 3). **B.** Oxidative degradation of 200 ng of DNA after 30 min incubation with 2 mM H_2_O_2_ and 50 μM FeSO_4_ in the absence or presence of 5, 10 or 25 mM polyP_300_ (in Pi-units), 75 mM NaH_2_PO_4_ or 75 mM KH_2_PO_4_. Samples were applied onto an agarose gel and visualized using ethidium bromide (n=3).

## Discussion

Iron serves as an essential co-factor in many enzymes, which are involved in processes ranging from metabolism to DNA synthesis and cell division. Yet, free iron is highly toxic under oxygen rich-condition as it readily undergoes the Fenton or Haber-Weiss reactions, thereby producing extremely reactive oxygen species, particularly hydroxyl radicals (Dixon & Stockwell, 2014) (Fig. 6A). Because of this dichotomy in cellular need for and risk of free iron, aerobically growing organisms such as *E. coli* tightly regulate their iron homeostasis. Here we demonstrate that cisplatin, an anti-cancer drug and broad-spectrum antimicrobial, triggers cytotoxicity in bacteria not only through its ability to crosslink DNA but also by causing cellular iron overload. By using a combination of genetic and biochemical tools, we discovered that bacteria defend themselves against this secondary insult through polyP, which effectively complexes the iron, prevents the Fenton reaction and mitigates cisplatin toxicity (Fig. 6B).

**Figure 6.**
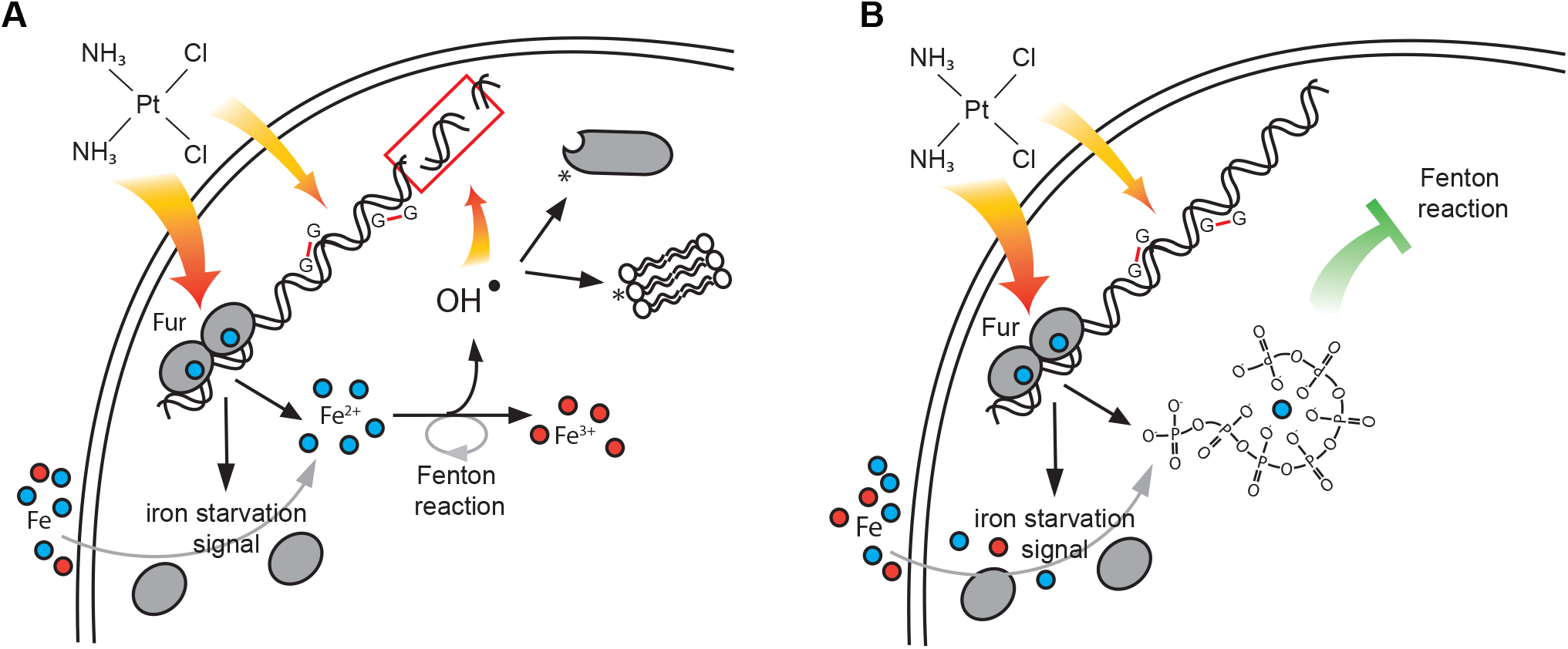
Model for the protective effects of polyphosphate against cisplatin. **A**. Our data suggest that cisplatin treatment causes iron accumulation that might contribute to a second mode of killing through Fenton-mediated oxidative stress and DNA damage. Iron stress likely results from a combination of events; the release of protein-bound iron, and the inactivation of the master repressor of bacterial iron homeostasis Fur that triggers gene expression changes aimed to increase the cellular iron load. Subsequently, cellular free iron levels increase and can wreak havoc under aerobic growth conditions by producing highly reactive oxygen species, such as hydroxyl radicals. **B**. Due to its polyanionic nature, endogenous polyP binds free iron, and prevents iron to act as a catalyst for the Fenton reaction, thereby drastically reducing the toxicity of cisplatin.

It is well established that polyP interacts with metals such as Ca^2+^ to form acidocalcisomes and protects organisms by complexing and sequestrating heavy metals (Docampo et al., 2005, Tocheva et al., 2013, Toso et al., 2011). In contrast, however, very little is known about the interaction of polyP with iron, the physiological relevance of such an interaction, or its potential role in the Fenton reaction (Rachmilovich-Calis et al., 2011, Richter & Fischer, 2006). A much-needed search for physiologically relevant Fenton inhibitors revealed that nucleotide phosphates, such as ATP, either stimulate or prevent the Fenton reaction, depending on the number of iron coordination sites that were occupied by phosphates in the complex (Richter & Fischer, 2006). The most effective Fenton inhibitor turned out to be dimers of ATP-y-S, which, through their six existing phosphate groups, block all iron coordination sites. This appears to prevent H_2_O_2_ from reacting with iron and mitigates hydroxyl radical formation. We now propose that polyP, due to its polyanionic nature and structural flexibility, is able to also occupy all relevant coordination sites in iron, thereby interfering with peroxide binding and preventing the Fenton reaction (Fig. 6B).

Our studies revealed noteworthy parallels between the protective function of polyP in bacteria exposed to cisplatin and in bacteria that lack the iron repressor Fur. Our transcriptional analysis of cisplatin-treated wild-type bacteria supported this conclusion by demonstrating that close to 80% of previously identified Fur-regulons are upregulated in bacteria treated with cisplatin (Seo et al., 2014). At first glance, these results suggested that bacteria treated with cisplatin suffer from stress conditions that trigger intracellular iron depletion; hence, the upregulation of siderophores and transport mechanisms aimed to replenish the intracellular iron pools. However, analysis of the cisplatin response in polyP-depleted bacteria yielded quite the opposite result, and, in fact, suggested that cisplatin triggers an accumulation of free iron, which becomes highly toxic and mutagenic unless chelated by polyP. Further support for this conclusion came from cisplatin-treatment studies in Fe-deplete media, as well as complementation studies in which we either overexpressed the *E. coli* Fe-storage protein FtnA, or *rhyB*, a small RNA that down-regulates the amount of Fe-S cluster proteins in bacteria. In all three scenarios, cisplatin-treatment of the *ppk* deletion strain no longer increased the mutagenesis rate or affected survival beyond what we observed in cisplatin-treated wild-type *E. coli.* These results strongly argue that cisplatin treatment causes an increase in intracellular iron, which, unless chelated by polyP, significantly increases the toxicity of cisplatin. Since cisplatin is well known for its ability to interact with and bind to cysteine and methionine residues in proteins (Peleg-Shulman et al., 2002), and Fur contains a redox sensitive cysteine-coordinating zinc site (d’Autréaux et al., 2007), we now speculate that Fur itself might become a target of cisplatin. Inactivation of Fur would misleadingly send an iron starvation signal to the cell, causing iron accumulation. Intriguingly, very recent studies showed similarly disruptive effects of cisplatin on the iron homeostasis in cancer cells (Miyazawa et al., 2019). In contrast to bacteria, however, which ultimately suffer from iron overload, cisplatin-treated cancer cells experience true iron starvation. This iron starvation phenotype is triggered by the covalent modification of two cysteines in the iron regulatory protein 2 (IRP2), a central activator of the mammalian iron starvation response (Miyazawa et al., 2019). Once inhibited, IRP2 is unable to downregulate the iron chelator ferritin, causing persistent iron depletion. Our realization that polyP serves as iron chelator helps to explain our recent finding that endogenous levels of polyP positively correlate with apoptosis in cisplatin-treated cancer cells (Xie et al., 2019). We now reason that by chelating iron, polyP further potentiates the iron starvation phenotype in cancer cells, hence causing the observed increase in cytotoxicity (Miyazawa et al., 2019, Xie et al., 2019).

The ability of polyP to complex iron might also contribute to its protective function under oxidative stress conditions, as well as other cellular insults that cause protein-bound iron to be released. Chelation of iron by polyP will not only prevent secondary oxidative stress conditions but would allow iron to be rapidly re-incorporated into proteins without the activation of complex and energy-consuming uptake systems once the stress is removed. It remains to be tested whether and how association of polyP with iron affects some of the other known functions of polyP, such as its ability to interact with and stabilize unfolding proteins, or its activity in blood coagulation and inflammation (Morrissey et al., 2012), adding a potentially new layer of complexity to this structurally simple molecule.

## Material and Methods

### Bacterial Strains and Growth Conditions

All strains, plasmids, and oligonucleotides used in this study are listed in Tables S5-S7, respectively. Gene deletions were generated by λ red-mediated site-specific recombination (Datsenko & Wanner, 2000). All chromosomal mutations were confirmed by PCR. *E. coli* MG1655 was grown at 37°C in lysogenic broth (LB; Fisher) or MOPS minimal medium (Teknova) containing 0.2% w/v glucose and 1.32 mM K_2_HPO_4_ (MOPS-G). To conduct cisplatin resistance tests on plates, M9 minimal medium (Sambrook et al., 1989) containing 0.2% glucose (M9-G) was supplemented with 1.5% agar. When indicated, 0.2% w/v gluconate was used instead of glucose (M9-gluconate). For iron-depleted conditions, M9-G medium was mixed and incubated with 2 g/l Chelex100 (BioRad) for 1 hr at room temperature under constant agitation. The chelated solution was then sterile-filtered. For pBAD expression induction, arabinose (0.2% w/v) was added 15 minutes prior to the experiment in case of liquid assays or added to the media in case of plate assays. The following antibiotics were added when appropriate: Chloramphenicol (30 μg/ml), rifampicin (200 μg/ml), kanamycin (50 μg/ml), or ampicillin (100 μg/ml).

### Growth under iron starvation

Iron starvation growth assays were performed as described in (Outten et al., 2004). Briefly, *E. coli* MG1655 wild-type, *ppk* and *ppx* deletion strains were grown overnight in M9-gluconate medium, diluted and cultivated at 37°C until OD_600_ of 0.5 was reached. Cells were then diluted 1:100 into M9-gluconate medium supplemented with the indicated concentrations of 2,2’ bipyridyl (stock dissolved in 100 mM DMSO; Sigma-Aldrich). Each condition was performed in triplicate and growth was monitored for 18h. The OD_600_ reached after 18h was then normalized to the corresponding growth rate in M9-gluconate in the absence of chelator and plotted.

### Cisplatin survival assay

*E. coli* MG1655 wild-type and the isogenic mutant strains were grown at 37°C with shaking in MOPS-G medium to an OD_600_ ~0.4–0.8 and harvested by centrifugation. To analyze cisplatin sensitivity in liquid culture, the cells were resuspended in MOPS-G medium to an OD_600_ of 0.4, supplemented with various concentrations of cisplatin (stock solution 0.9 mg/ml in sterile _dd_H2O; Sigma-Aldrich) and incubated at 37°C with shaking (200 rpm). At defined time points of incubation (1 h – 20 h), the cells were harvested by centrifugation, washed twice, 10-fold serially diluted and plated onto LB agar. Survival was assessed after overnight incubation at 37°C. To determine the survival of bacteria when grown on cisplatin-containing plates, bacteria were cultivated in MOPS-G media until OD_600_ of 0.5 was reached. Then, the bacteria were 10-fold serially diluted and plated onto M9-G plates containing various concentrations of cisplatin. Colonies were counted after a 24h incubation at 37°C. To compare the cisplatin sensitivity of bacteria in iron-deplete liquid media, the bacteria were first grown in MOPS-G media as described above. Once an OD_600_ of 0.4 was reached, the bacteria were centrifuged, washed and resuspended in either untreated M9-G (+Fe) media or chelexed M9-G (-Fe) media in the absence or presence of 10 μg/ml of cisplatin. At defined time points of incubation (1h – 20h), the cells were harvested by centrifugation, washed twice, 10-fold serially diluted and plated onto LB agar.

### Mutagenesis assay

Mutagenesis rates were measured as described (Krisko & Radman, 2013). Briefly, cells were grown overnight in MOPS-G medium, diluted 1:100 into 30 ml fresh MOPS-G medium and cultivated at 37°C until OD_600_ of 0.5 was reached. The bacterial suspension was either left untreated or supplemented with 10 μg/ml cisplatin, and further incubated for 1hr. After the incubation, untreated and treated bacteria were washed twice in MOPS-G media and resuspended in 5 ml of LB media. The bacteria were then incubated overnight at 37°C with shaking. Serial dilutions were made and plated onto both LB agar and LB-rifampicin agar plates. After 24 h of incubation at 37°C, the colony forming units (CFU) were scored. The mutation frequency was calculated by dividing the number of rifampicin-positive colonies by the total number of colonies.

### RNAseq analysis

Four biological replicates of wild-type *E. coli* MG1655 and *Δppk* were cultivated in MOPS-G medium at 37°C until OD_600_ of 0.4–0.5 was reached. Cells (1 ml) were harvested either before or 15 min after the treatment with 20 μg/ml cisplatin in 1 ml of ice-cold methanol (– 80°C) to stop transcription. After centrifugation and removal of the supernatant, total RNA was prepared using the Ambion RiboPure-Bacteria Kit (Thermo Fisher Scientific) according to the manufacturer’s instructions. The samples were DNase I treated, followed by depletion of rRNA using the Illumina Ribo Zero Kit (Illumina) for Gram-negative bacteria. Fifty base single end sequencing was performed on an Illumina HiSeq 4000 using the University of Michigan DNA Sequencing Core. Sequence reads from the RNA-seq were mapped onto the reference genome (NC_000913). Genes with a log2 fold change >0.5 and an FDR value <0.01 were considered as differentially expressed genes (DEGs).

### GOC enrichment and analysis

Differentially regulated genes upon cisplatin treatment and/or between strains were categorized according to their annotated COG categories (Ashburner et al., 2000). Functional enrichment of COG categories was determined by performing a modified one-tailed Fisher’s Exact test (EASE score from DAVID), with a *p*-value <0.05 considered significant. Comparison to iron-regulated genes (Seo et al., 2014) seen in Venn Diagram in Fig. 2C, was performed using a Fisher’s Exact test. The heatmap was produced using open-source statistical software R (https://www.r-project.org/) with log2 fold change data. Figures 2 and S2 were also produced in R.

### Enzymatic assays

The aconitase activity assay was performed according to the manufacturer’s protocol (MAK051, Sigma Aldrich). Briefly, bacterial cell pellets were resuspended in 1 ml of aconitase lysis buffer (0.1 mM Tris-HCl, pH 8.0, 0.1 M KCl, 1 mM PMSF, and 0.6 μg/ul lysozyme) and lysed by 5-6 rounds of freeze & thaw cycles. The lysates were centrifuged at 14,000 rpm for 10 min at 4°C. The total protein concentration was determined using the Bradford assay. The aconitase assay was performed by mixing 100 μg of cell lysate with 200 μl of 1x aconitase assay buffer (0.6 mM MnCl_2_, 25 mM sodium citrate, 0.25 mm ADP, 50 mM Tris-HCl, pH 7.6). The absorbance was recorded at 340 nm, and the specific aconitase activity was calculated per milligram of total proteins (Kaur et al., 2017).

### *In vitro* Fenton reaction and DNA damage assay

The ABTS assay was performed as previously described in (Zheng & Huang, 2014). In brief, 250 μM ABTS was mixed with 5 μM FeSO_4_ and 200 μM H_2_O_2_ in the absence or presence of increasing concentrations of polyP^300^ (see figure legends for details). The assay was performed in 10 mM acetate buffer, pH 3.6 to control acidity. After 30 min of incubation at 37°C, the mixture was read at an absorbance of 414 nm. The DNA damage assay was performed by mixing 50 μM FeSO_4,_ 5 mM H_2_O_2_ and 10 μM linearized DNA from the plasmid pBAD18 in water with the indicated concentration of either Na_2_HPO_4_, KH_2_PO_4_, or polyP_300_ (concentration given in P_i_-units). The reaction was initiated by the addition of H_2_O_2_ and incubated for 30 min at 37°C. Then, samples were loaded onto an agarose gel. Staining with ethidium bromide was used to visualize the DNA bands.

## Acknowledgements

We thank the DNA Sequencing Core (BRCF), the Bioinformatics Core of University of Michigan and Christopher Sifuentes for RNA sequencing and data analysis. We thank Jan Dahl, Michael Gray and the entire Jakob lab for helpful discussions and important input. This work was supported by NIH grants GM122506 to U.J., a NIH T32 Career Training in the Biology of Aging grant to E.Q.; F.B. was funded by an EMBO long-term fellowship (ALTF 601-2016).

